# Region-specific impact of aging on cortical myelination and thickness

**DOI:** 10.1101/2023.02.04.527112

**Authors:** Arianna Brancaccio, Davide Tabarelli, Paolo Belardinelli

**Affiliations:** Center for Mind/Brain Sciences—CIMeC, University of Trento, I-38123 Trento, Italy

**Keywords:** brain aging, cortical development, myelo-architecture, cortical thickness, magnetization transfer imaging, T1w/T2w ratio imaging

## Abstract

Healthy aging affects both grey and white matter. However, the trajectories of regional specific degeneration are not fully understood. Here we investigate the effects of aging on cortical thickness and myelin concentration in a large cohort of healthy participants (N = 610) aged between 18 and 89 years’ old who underwent single-site T1-weighted, T2-weighted and MTI sequences in the context of the Cam-CAN project. Participants were subdivided in three age groups representative of young, middle and late adulthood. The large size of the dataset allowed us to minimize the impact of sample variance without relying on multi-site acquisition protocols. We assessed linear changes in cortical thickness and cortical myelin concentration; the latter was assessed using both T1w/T2w ratio and MTR proxies, to evaluate which is the most stable metrics. Our results do not fit with either the anterior-posterior gradient or the last-in/first-out hypothesis. We demonstrate that aging patterns are more complex than just depending on a spatial gradient or the temporally reversed order of regional development. Moreover, we show a dissociation in aging patterns between somatosensory and motor regions both in terms of cortical thickness and myelin concentration. Finally, comparing T1w/T2w and MTR results of cortical myelination, we found the latter being a more stable and reliable proxy.

**Highlights:**

- Cortical thickness and myelo-architecture changes must be jointly considered in investigating brain aging trajectories.
- We assessed linear changes in cortical thickness and myelination in a large, homogeneous and single site MRI dataset.
- Motor and sensory regions show a dissociation in their aging trajectories both in terms of cortical thickness and myelin concentration.
- Sensory processing regions show similar aging trajectories in both cortical thickness and myelin concentration.
- MTR is a more reliable proxy for myelin concentration compared to T1w/T2w ratio.

## Introduction

An extended consensus has converged on general age-related cortical degeneration due to grey and white matter thinning (Salat et al., 2004; Raz, 2000; Gunning-Dixon et al., 2008; Davis et al., 2009). The majority of previous works have focused on grey matter changes. However, two relevant issues need to be stressed: first, most studies on cortical thickness changes associated with healthy aging did not rely on large samples representative of the full life span. Secondly, cortical thinning is not the only type of age-related structural brain change: cortical loss of myelin concentration in the cortex should be taken into account among other factors.

As for the first point, most of previous studies have investigated cortical thickness changes due to aging including small sample sizes. For example, a sample size of approximatively N = ~75 (Salat et al, 2004; Thambisetty et al., 2010; Hutton et al., 2009), even if sufficiently large with respect to a typical non age-related structural study, may still be insufficient to be sensitive to the small age-related cortical degenerations effects (Fjell et al., 2009). Moreover, several studies suffer from nonuniform or limited age distribution in the sample and/or multi-site analysis pipelines. Both consistent and divergent results in terms of age-related regional-specific thinning have been reported in these studies. Frontal cortices consistently appear the most affected by aging (Salat et al, 2004; Thambisetty et al., 2010; Hutton et al., 2009). Differently, discrepancies have been found in the age-related changes especially in parietal regions, with some studies claiming that thinning rate in this region is sometimes larger compared to frontal areas’, while other works found the opposite pattern (e.g., Salat et al., 2004; Thambisetty et al., 2010; Hutton et al., 2009). The reasons behind discrepancies in previous results have been discussed in a review by Raz and Rodrigue (2006) and attributed to the variability due to heterogeneity in the samples, small sample sizes and different analysis pipelines employed. Recent studies have made attempts to overcome these issues (e.g., Fjell et al., 2009; McGinnis et al., 2011; Lemaitre et al., 2012; Frangou et al., 2020). These works increased the number of participants per age group or used multiple samples to test the reproducibility of age-related effects. However, the sample sizes employed might have been still not sufficiently sensitive to age-related changes in the brain structure, as sample heterogeneity cannot be controlled for in non-longitudinal settings. Also, to increase the number of participants, multi-site recordings have been included, in principle further increasing sample variability (See Suppl. Table 1 for a list of studies and their sample characteristics). As a matter of fact, both consistency and discrepancy emerged in these latter results, with evidence both in favour and in contrast with the *last in first out* hypothesis (Raz, 2000), according to which areas that develop late (e.g. frontal) also degenerate early. In this regard, it is worth noticing that the highest variability in terms of age-related cortical thinning has been found in frontal regions (Frangou et al., 2020). Therefore, attempts at describing age-related brain changes in participants covering broad age-ranges should employ large, single-site datasets in order to optimally extract small age-related structural effects from the sample variance, inevitably contaminated by heterogeneity and inter-individual variability.

Cortical thickness alterations do not represent the only degeneration process related to aging. One further relevant biomarker of aging is cortical myelin concentration. However, also in this case, evidences are based on relatively small sample sizes and different metrics, with eterogeneous properties and drawbacks. In this regard, it is important to describe the key differences between the T1w/Tw ratio and MTR. A direct MR measure of myelin content in brain tissues (i.e. by exploiting the relaxation times of protons bound in myelin macromolecules) is impossible because these relaxation times are in the order of a few microseconds (Sled, 2018). Glasser and Van Essen (2011) conceived T1w/T2w ratio as an MRI-proxy sensitive to myelin content. This measure is based on the ratio between T1- and a T2-weighted intensity values and returns a unitless proxy of myelin tissue concentration (Shafee et al, 2015). However, not only myelin but also iron molecules contribute to the ratio of the T1w/T2w contrast (Shafee et al., 2015; Fukunaga et al., 2010. Since iron is accumulated over the life-span (Dröge et al., 2007), this represents a confound in investigating age-related myelin changes. One further limitation of this metric consists in the non-linear difference of intensity histograms of the images used to compute the ratio: a within-subject normalization procedure is necessary before T1w/T2w computation, a step potentially increasing inter-subject variability and in turn overall variance in group analyses. Moreover, the unitless quantity resulting from the T1w/T2w ratio after each individual histogram normalization, is still subject-dependent and therefore a group level z-value based normalization, assuming Gaussian statistics, has to be performed. Therefore, results on cortical myelin content might be affected not only by the effect of age-related iron accumulation, but also by non-optimal MRI images’ intensity normalization procedures within and across participants. Differently from the T1w/T2w ratio, the MTR is specifically designed to assess myelin content. MTI is based on magnetization transfer between populations of protons in brain tissues. Shortly, a saturation pulse is delivered to induce a magnetization in the pool of protons bound in macromolecules. Right after the saturation pulse, a standard sequence is used to probe relaxation times of free protons. This leads to a “saturated image”. The same procedure is repeated without the saturation pulse, leading to a “baseline image”. When the saturation pulse is applied, protons bound in macromolecules lose their magnetization by means of different relaxation processes. Among those, they partially transfer their magnetization to the free proton population. This leads to reduced intensity of T1w or T2w contrasts in the image acquired right after the saturation pulse. The MTR is then defined as the ratio between the intensity reduction and the baseline image. This unitless proxy ranging by design between 0 and 1 is proportional to the proton concentration bound in myelin macromolecules in the voxel of interest. Differently from T1w/T2w ratio, the MTR approach compares images acquired with the same parameter (free proton relaxation time). Therefore, no preliminary intensity normalization is required on the baseline and saturated images. In addition, since the MTR proxy is a unitless ratio bound by design between 0 and 1, it does not require group-level normalization. MTR is currently the most optimized non-invasive MR based measure of myelin content in the brain and it has also been shown to correlate better than the T1w/T2w ratio with histologic studies (van der Weijden et al., 2021; Hagiwara et al., 2018).

Highest myelination is consistently found in parietal regions in studies employing the T1w/T2w ratio (e.g., Glasser and Van Essen, 2011; Shafee et al., 2015; Kwon et al., 2020). However, discrepancies can be found in previous literature regarding the timing at which cortical myelin content develops in the different brain regions. Some works have proposed that sensory areas are the first to develop while the frontal cortices develop last, in accordance with the *last in first out* hypothesis (e.g., Deoni et al., 2015). Differently, other studies including participants from broad age ranges showed primary motor cortex with the highest increase in myelination during adolescence, young adulthood and in some cases, middle adulthood (e.g. Shafee et al., 2015; Kwon et al., 2020; Grydeland et al., 2013). Importantly, all the works cited above employed the so-called T1w/T2w ratio to assess cortical myelin content. Previous works employing the MTR proxy to assess age-related myelin changes, also reported contrasting evidence with parietal regions showing age-related myelin changes only in one out of two studies (Wu et al., 2016; Karolis et al., 2019). In fact, contrasting evidence has been found especially on the age-related myelination changes in parietal areas, both using T1w/T2w ratio and MTR. These differences in recent studies investigating more comparable and extended age span, might still be ascribed to limited sample sizes and different metrics employed (See Suppl. Table 2 for a list of studies, sample characteristics and metric employed), For this reason, a systematic investigation of the results obtained with T1w/T2w and MTR on the same samples needs to be accomplished to understand which metric is more stable in assessing myelin content over the lifespan.

Here, we attempt to overcome previous works’ limitations in terms of sample size and metrics employed. In fact, we investigate age-related changes both in gray matter and cortical myelin content in a large cohort of participants (N= 610; age: 18 – 89; Cam-CAN Dataset (Shafto et al., 2014; Taylor et al., 2017)). We assess cortical myelin content using both the T1w/T2w ratio and the MTR to assess the most stable and reliable metric of cortical myelin. In this work, we show different magnitudes of cortical thinning depending on the brain region. In particular, we report results not fittin with either the *anterior-posterior gradient* (Thambisetty et al., 2010) or *last in first out* (Raz, 2000) accounts. Moreover, we show MTR as a more stable metric of cortical myelin content compared to the T1w/T2w ratio. In fact, MTR results are less affected by confounding effects due to cortical thickness. Furthermore, we report a dissociation in the aging patterns of sensory and motor sectors, as these regions show different patterns of development from young adulthood to old age both in terms of grey and white matter thinning. Finally, we demonstrate that the relationship between the T1w/T2w ratio and the MTR is variable over the cortex, clearly suggesting the need of further investigating the relationship between the two metrics as well as the quantities they really measure.

## Methods

### Dataset and Acquisition

Data used in this work were obtained from the CamCAN repository (available at http://www.mrc-cbu.cam.ac.uk/datasets/camcan/), (Cam-CAN; Shafto et al., 2014; Taylor et al., 2017). Data were collected in compliance with the Helsinki Declaration (Cambridgeshire 2 Research Ethics Committee; reference: 10/H0308/50). In this work, MRI data from 610 healthy participants out of the approximatively 700 participants recruited at the Cambridge Centre for Ageing and Neuroscience (Cam-CAN; Shafto et al., 2014; Taylor et al., 2017) were included in this study. The Cam-CAN dataset is a large repository including both raw and preprocessed structural (T1w, T2w, DWI, MTR) and functional (sensorimotor task, movie-watching and resting state) MRI and MEG (sensorimotor task and resting state) data from healthy participants covering the whole adult lifespan. Subjects who underwent all the MRI scans described in the following Recording Section (T1w, T2w and MTI) were included in this study. This led to a final dataset of 610 healthy subjects (310 females) with age ranging between 18 and 89 years.

### MRI Recordings

All MRI datasets were collected in a single site (Cambridge Centre for Ageing and Neuroscience) using the same 3 T Siemens TIM Trio scanner with a 32-channel head coil. Each participant underwent: a T1-weighted Magnetization Prepared Rapid Gradient Echo (MPRAGE) sequence with repetition time (TR) = 2250 ms; echo time (TE) = 2.99 ms; inversion time (TI) = 900 ms; flip angle = 9°; field of view (FOV) = 256 × 240 × 192 mm; resolution = 1 mm isotropic;); a T2-weighted single slab three-dimensional turbo spin echo SPACE sequence (Mugler and Brookeman, 2004) with TR = 2800 ms; TE = 4.08 ms; flip angle α = 9°; field of view (FOV) = 256 × 240 × 192 mm; resolution = 1 mm isotropic. To evaluate myelin content, each participant underwent a Magnetization Transfer Imaging session (MTI) consisting of two MT-prepared Spoiled Gradient (SPGR) sequences, with and without the application of a Gaussian shaped RF pulse (offset frequency 1950 Hz, bandwidth 375 Hz, duration 9984 μs) for myelin protons saturation. The SPGR sequence parameters were TE = 5 ms; flip angle α = 12°; field of view (FOV) = 192 × 192 mm; resolution =1.5 x 1.5 mm; the repetition time were set to 50 or 30 ms, depending on the estimated SAR for the subject exceeding or not the limits, respectively. For more detailed information on the MRI recordings parameters collected during Stage 2 of the Cam-CAN project, the reader is invited to refer to Taylor et al. (2017).

### Data Preprocessing

T1-weighted images were first processed to define anatomy and extract cortical thickness values over the cortex. Each volume was denoised and debiased for the correction of field inhomogeneities using SPM software (Friston, 2003). Then, the Computational Anatomy Toolbox 12 (CAT12) (Gaser et al., 2022) was used for the segmentation of Gray Matter (GM), White Matter (WM) and Cerebro Spinal Fluid (CSF). Also, the total intracranial volume (TIV) was computed and the brain extraction implemented. Moreover, each volume was non-linearly registered on a template in MNI space (IXI database of 555 subjects; http://www.brain-development.org). The method implemented in CAT12 was then used to extract the cortical thickness values while concurrently defining the mid-thickness central surface as a mesh of approximatively 327000 vertices (Dahnke et al., 2013). The mid-thickness surface is anatomically defined as the surface halfway between the pial and white/gray matter interface. A final spherical mapping-based projection onto the FreeSurfer average template (with a Gaussian smoothing kernel of FWHM = 15 mm) was performed in order to allow for group analyses (Fischl et al., 2012). T2-weighted images were also denoised and debiased for the correction of field inhomogeneities using SPM. Then, they were co-registered (rigid body co-registration) on the individual T1-weighted image. After these steps, both T1- and T2-weighted images were calibrated using a non-linear approach (Ganzetti et al., 2014) based on the intensity extracted from internal brain tissues (GM, WM, CSF) (Glasser and Van Essen., 2011). At this point, the ratio between the T1- and T2-weighted images was computed. The final volumetric values of T1w/T2w ratio were then extracted on the mid-thickness mesh, resulting in a surface-based map of myelin content as estimated by T1w/T2w ratio. A final spherical mapping projection onto the FreeSurfer average template (with a Gaussian smoothing kernel of FWHM = 5 mm) was performed in order to perform group analyses. Also, the MTI images (with and without saturation pulse, denoted as M_0_ and M_S_ respectively) were co-registered to the T1-weighted (rigid body co-registration) and the brain extracted using the brain mask previously computed in the T1w analysis. Successively, the MT ratio was computed as (M_0_–M_S_)/M_0_ (Wolff and Balaban, 1989; Mehta et al., 1996). Finally, the volumetric MTR values were extracted on the mid-thickness mesh, resulting in a surface-based map of myelin content as estimated by MTR. Finally, a spherical mapping projection on to the FreeSurfer average template (Gaussian smoothing kernel of FWHM = 5 mm) was performed to allow group analyses on MTR as well. At the end, we obtained a total of 610 mid-thickness surface maps of thickness, T1w/T2w ratio and MTR projected on the FreeSurfer average group template.

### Data Analysis and Statistics

Before computing group analyses, the following steps have been performed. First, maps of thickness, T1w/T2w ratio and MTR were reduced from the original 327000 vertices to 7117 regions (henceforth named *isorois)* defined by pooling together 4^th^ order neighbors on the mesh. Furthermore, the MTR maps were corrected for the different repetition times (TRs) used in the acquisition. In fact, MTR sequences had been recorded with two different TRs depending on participants’ SAR estimation (TR = 50 ms or TR = 30 ms). TR influences the temporal distance between the saturation pulse and the following standard Spoiled Gradient (SPGR) sequence. Smaller TR (SPGR sequence closer to saturation pulse) results in overall greater ratio for myelin content. Therefore, prior to pooling all MTR results together for group analysis, the following correction was applied: the mean of the distributions of MTR surface values (all subjects with TR = 50 ms or TR = 30 ms) was computed; since the two distributions were well gaussian shaped (see Suppl. Fig. 6), all TR = 50 ms MTR data were corrected by subtracting the mean difference with respect to the TR = 30 ms dataset. Finally, the T1w/T2w ratio maps were z-values corrected for each subject, assuming the distribution of all T1w/T2w ratio values on the whole subject mid-thickness surface being a single Gaussian distribution. At this point, data were ready for group analyses. In order to implement them, subjects were divided in three age groups (from 18 to 90 y.o in 24 y steps; age ranges 18-42/43-66/ 67-90; N = 185/212/190). All dipoles (from all subjects belonging to a given age group) of each isoroi were pooled together (separately for each age group) and the average values of CT, T1w/T2w z-ratio corrected and MTR (TR corrected) were computed. For the sake of clarity, the TIV was regressed out as a confound for each quantity (CT, T1w/T2w and MTR) and from now on, CT, Tw1/T2w and MTR will be always intended as TIV-corrected. Furthermore, the linear change with age was assessed by means of a linear fit for each quantity and isoroi. To understand how myelin content influences the cortex thinning, we further regressed out from CT both our proxies of myelin content (T1w/T2w ratio and MTR, separately). This resulted in two additional thinning maps (slopes of the linear fits on the cortex) representing how CT changes with age when either the T1w/T2w ratio or the MTR were regressed out as a confound. The same approach was used to conversely understand how myelin content is affected by cortical thinning, resulting in two further surface maps with the slopes of myelin data linear fits where the CT was regressed out from the T1w/T2w and MTR. Furthermore, two additional linear fits between CT and T1w/T2w ratio and MTR were computed to understand the direct relationship between CT and myelin content changes in the three different age groups. The last comparison was performed to estimate the consistency of the two proxies used for myelin content assessment. All results from the above linear fits were thresholded at p < 0.05 and corrected for FDR with q = 0.05, considering the total number of isorois (7117). Furthermore, a quantitative analysis was implemented on each of the metrics’ averages and linear fits’ slopes described to appreciate the aforementioned changes. Average values and significant slopes (linear fit p < 0.05 with respect of a null-hypothesis of a flat line) were extracted for all the isorois belonging to the frontal, sensory, motor, premotor and occipital sectors separately. We defined these sectors using ROIs from the HCP multimodal atlas (Glasser et al., 2016). For a list of ROIs included in the different sectors see Table 1. These values, pooled by sector, were then statistically compared across the three age groups with a two-tailed non-paired t-test, in order to exploit significant differences at a whole sector level.

**Table 1:**
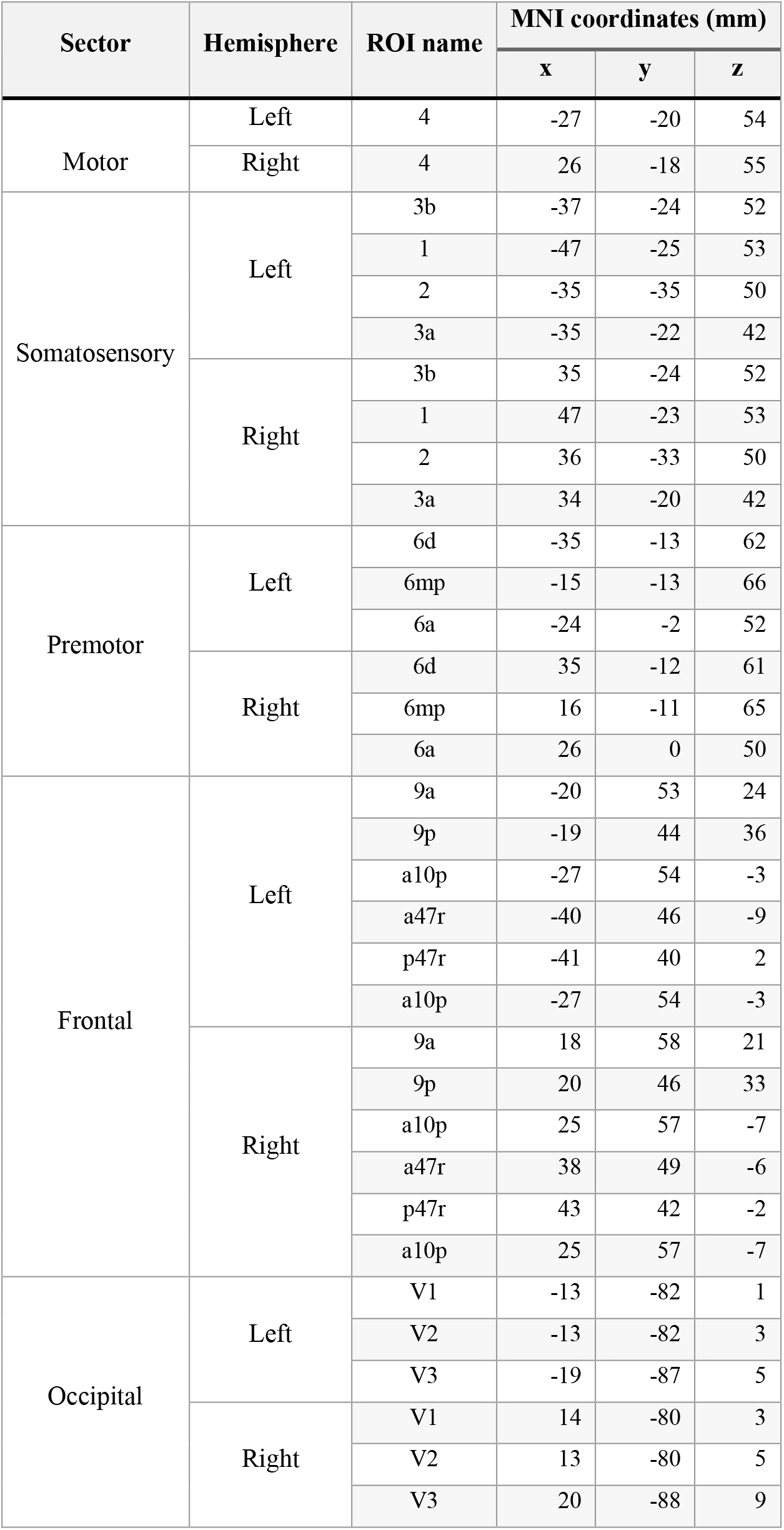
List of ROIs from the HCP multimodal atlas included in each sector of interest. For each ROI, the MNI coordinates are reported.

| Sector | Hemisphere | ROI name | MNI coordinates (mm) |  |  |
| --- | --- | --- | --- | --- | --- |
|  |  |  | x | y | z |
| Motor | Left | 4 | -27 | -20 | 54 |
|  | Right | 4 | 26 | -18 | 55 |
| Somatosensory | Left | 3b | -37 | -24 | 52 |
|  |  | 1 | -47 | -25 | 53 |
|  |  | 2 | -35 | -35 | 50 |
|  |  | 3a | -35 | -22 | 42 |
|  | Right | 3b | 35 | -24 | 52 |
|  |  | 1 | 47 | -23 | 53 |
|  |  | 2 | 36 | -33 | 50 |
|  |  | 3a | 34 | -20 | 42 |
| Premotor | Left | 6d | -35 | -13 | 62 |
|  |  | 6mp | -15 | -13 | 66 |
|  |  | 6a | -24 | -2 | 52 |
|  | Right | 6d | 35 | -12 | 61 |
|  |  | 6mp | 16 | -11 | 65 |
|  |  | 6a | 26 | 0 | 50 |
| Frontal | Left | 9a | -20 | 53 | 24 |
|  |  | 9p | -19 | 44 | 36 |
|  |  | a10p | -27 | 54 | -3 |
|  |  | a47r | -40 | 46 | -9 |
|  |  | p47r | -41 | 40 | 2 |
|  |  | a10p | -27 | 54 | -3 |
|  | Right | 9a | 18 | 58 | 21 |
|  |  | 9p | 20 | 46 | 33 |
|  |  | a10p | 25 | 57 | -7 |
|  |  | a47r | 38 | 49 | -6 |
|  |  | p47r | 43 | 42 | -2 |
|  |  | a10p | 25 | 57 | -7 |
| Occipital | Left | V1 | -13 | -82 | 1 |
|  |  | V2 | -13 | -82 | 3 |
|  |  | V3 | -19 | -87 | 5 |
|  | Right | V1 | 14 | -80 | 3 |
|  |  | V2 | 13 | -80 | 5 |
|  |  | V3 | 20 | -88 | 9 |

## Results

### Age-related changes in Cortical Thickness

First, the CT changes with age and their relationship with cortical myelin content are reported. As it can be appreciated from Figure 1A, the CT average values of depend on the age group. In fact, in the early adulthood group (18 – 42 y.o.), the lowest values of CT are registered in sensory and occipital cortices (~ 2 mm), with frontal and premotor parietal areas showing larger CT values (~ 3 mm). Differently, in the mid-adulthood group (42 – 66 y.o.) and late adulthood group (66 – 89 y.o.), the CT is generally reduced (~ 2 mm), with anterior regions (frontal, premotor and motor) showing larger thickness than posterior ones. The changes in the sectors of interest can be better appreciated in Figure 1B, where the results from t-tests between age groups on the average CT are shown, together with violin plots describing the shape of CT values’ distribution in each brain sector of interest and for each age group, separately. From these quantitative analyses, all sectors show a decrease in CT average values with age, with all contrasts being significant (p < 0.0001). Confirming the results in Figure 1A, also in Figure 1B the motor, sensory and occipital cortices show the lowest CT average values compared to frontal and premotor cortices and these regions show largest CT also in the third group, despite CT loss with age.

**Figure 1:**
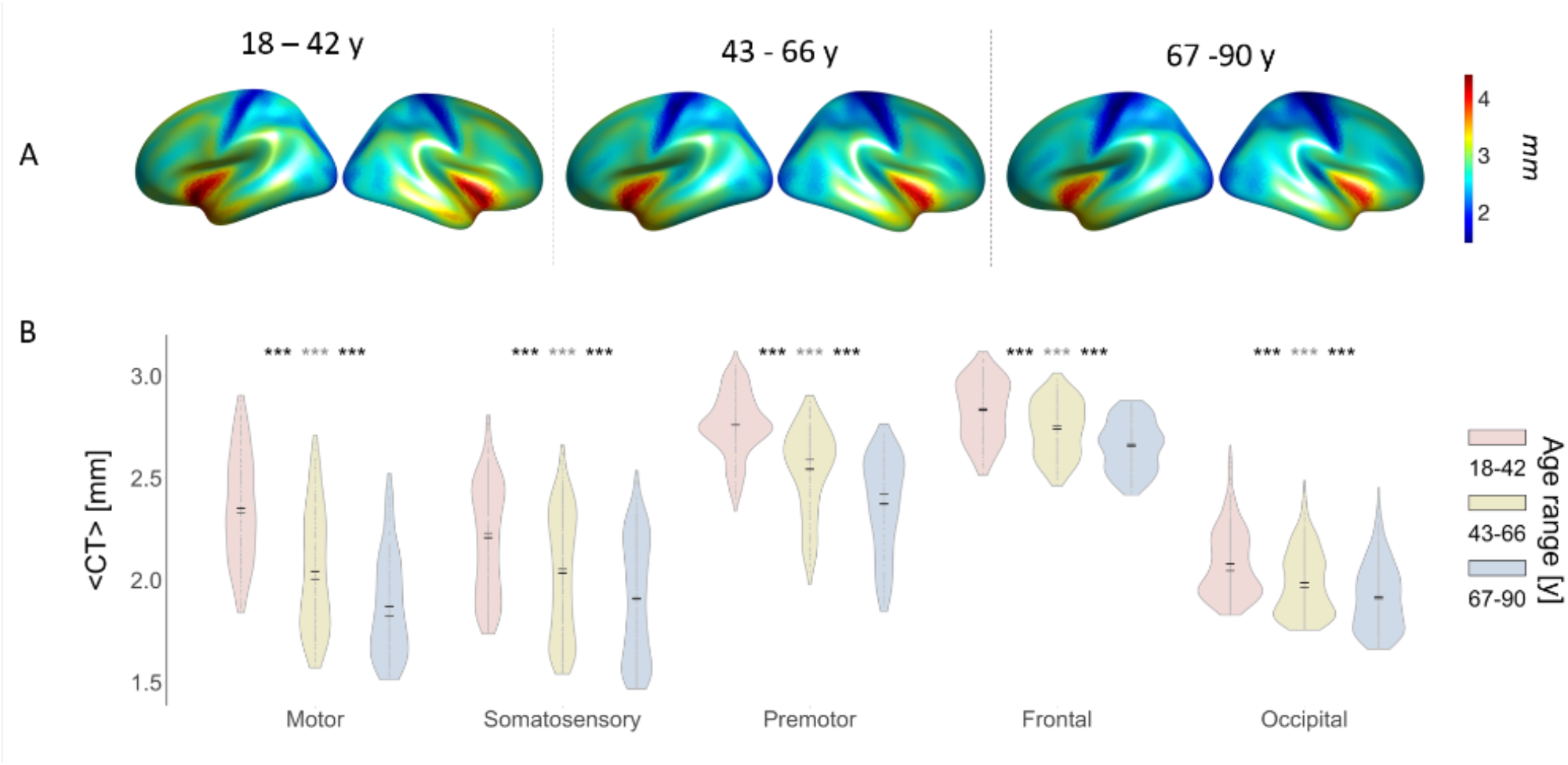
Results for average cortical thickness (CT) in the three age groups; A) CT average maps for the 7117 isorois for the three age groups separately; B) CT values distribution for each group in each brain sector of interest (mean in black, median in grey); significance asterisks summarize the results of a t-test between the three age groups for each brain sector separately (*** for p< 0.0001; ** for p < 0.001; * for p< 0.05). Grey asterisks refer to the comparison between the first and the third group.

In Figure 2A.1, the CT slope is reported separately for the three age groups. From the linear fits between CT and age, the overall trend is a loss between 0.01 and 0.02 mm per year in the young adulthood group, with the motor and premotor cortex showing the largest thinning. Differently, in the second mid-adulthood group, the CT slope tends to stabilize especially in the premotor and motor cortex. In the third late adulthood group, the linear fit between CT and age shows a greater thinning compared to the second age group in frontal and occipital areas. In general, the somatosensory cortex appears to be the most stable cortical area over the age groups. Figure 2B.1 shows the distribution of the CT slope for each group and the results from the t-tests across the age groups, separately for the different brain regions of interest. The decreased slope in sensory cortex thinning is confirmed by the sector-based analyses: the sensory cortex shows a significant thinning rate in the middle-age group compared to the young age group and a significant difference in CT slope between the first group and the third one (p < 0.0001), but no significant difference in CT slope values between second and third group. Differently, in the motor and premotor cortex the thinning rate is fast in the first group and then gets significantly slower with age (for all the contrasts p < 0.0001, except for the premotor cortex contrast between the first and second group which shows no differences in CT slope). Therefore, sensory cortices show a stable thinning over the life span, while premotor and motor cortices suffer from a fast thinning in the first age group which slows down with aging. Differently from sensory and pre-/motor districts, frontal and occipital cortices show similar behaviors, with the thinning rate getting significantly faster as age increases (for all contrasts p < 0.0001).

**Figure 2:**
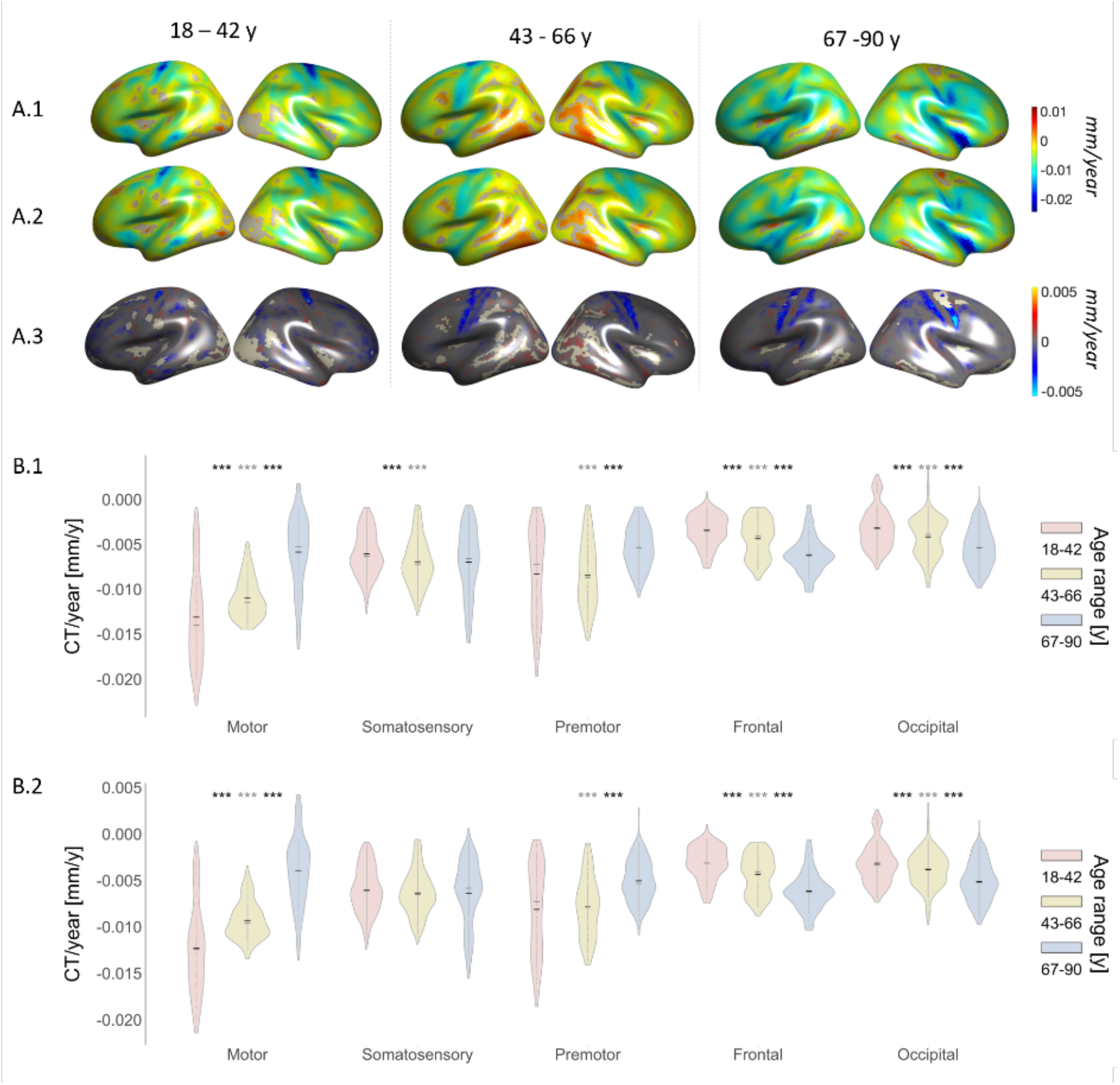
Results for the linear fit between Cortical Thickness and Age (Cortical thinning/CT Slope) in the three age groups (with and without MTR regressed out from CT before computing the linear fit between CT and Age); A.1) Source map of the slope values from the linear fit between CT and Age; A.2) Source maps of the slope values from the linear fit between CT and Age, with MTR regressed out as a confound.; A.3) Source map of the difference between A.1 and A.2: the difference is computed only between the values that are significant in both original contrasts; B.1) CT slope values distribution for each group in each brain sector of interest (mean in black, median in grey); B.2) CT slope values distribution with MTR regressed out as a confound for each group in each brain sector of interest (mean in black, median in grey); significance asterisks summarize the results of a t-test between the three age groups for each brain sector separately (*** for p< 0.0001; ** for p < 0.001; * for p< 0.05). Grey asterisks refer to the comparison between the first and the third group.

As for the relationship between CT slope and myelin, in Figure 2A.2 we report the results for the linear fits between CT (with MTR regressed out) and age to investigate how the relationship between CT and age changes when the effects of the myelin content (as assessed with MTR) are not taken into account. To better appreciate the differences between Figure 2A.1 (CT vs age) and 2A.2 (CT with MTR regressed out vs age), we computed the contrast reported in figure 2A.3, representing the difference between figure 2A.1 and 2A.2 (the difference is shown only for values which are significant in both original surface maps). In Figure 2A.3, a visible difference due to MTR effects in the linear fit between CT and age is shown. In this regard, Figure 2A.3 shows the main effect of MTR on the relationship CT-age. This is visible over the parietal district in all age groups (particularly in the second and third age groups). A slight increase in the age-related thinning rate (faster CT slope) of ~ 0.005 mm per year over the parietal cortex is detectable when the MTR is regressed out from CT. In figure 2B.2 (distribution of the CT slope values when the MTR is regressed out) the sensory cortex appears the most affected area by MTR regression. In fact, the CT slope differences in the sensory cortex become non-significant across age groups when the MTR is regressed out from the CT slope. Frontal, premotor, motor and occipital regions do not suffer from MTR regression in terms of CT slope, as their result remains the same as in figure 2B.1, when the MTR effect on CT is not regressed out from the CT before computing the linear fit with age.

The same logic was applied in the investigation of the relationship between CT and age, depending on whether the contribution of the myelin content as assessed with the T1w/T2w ratio this time, is taken into account or not. In this regard, Suppl. Figure 1A.1 reports the results for the linear fit between CT and age, while Suppl. Figure 1A.2 reports the result for the contrast between the CT and age when T1w/T2w is regressed out from CT (CT slope when T1w/T2w ratio is regressed out). Also here, the difference between Suppl. Figure 1A.1 and Suppl. Figure 1A.2 becomes remarkable computing the difference between the significant values obtained from the two linear fits between CT and age (when T1w/T2w ratio is regressed out). This difference is reported in Suppl. Figure 1A.3, where myelin content estimation with T1w/T2w ratio does not provide for detectable group difference in the relationship between CT and age as in case of myelin content estimated with the MTR. In fact, by comparing Figure 2A.3 and Suppl. Figure 1A.3, the cortical myelin content appears to increase the thinning rate when this is estimated with MTR and regressed out from CT in its linear fit with age. On the contrary, the same does not seem to happen when the myelin content is estimated with T1w/T2w ratio. In this regard, comparing Suppl. Figure 1B.1 and Suppl. Figure 2B.2, it can be appreciated that the CT slope does not change in any region’s contrast between groups when the T1w/T2w ratio effect is regressed out from the linear fit between CT and age.

### Age-related changes in Cortical Myelin

The same analyses (average, linear fits and sector-specific t-tests between age groups) were run on the cortical myelin content proxies. As for the average MTR, in figure 3A, first age group, the frontal regions are the least myelinated and myelination levels in these areas tend to be larger in the second group compared to the first, but then to decrease again in the third group, suggesting a tendency towards an inverted U-shaped trajectory of MTR. On the other hand, the sensory, motor and occipital districts appear as the most affected by age in terms of myelin content. In these regions, the average MTR decreases dramatically from first to the third group. The premotor district appears more stable in terms of myelin content since changes in these regions become clear only in the third group. These results are further corroborated by quantitative analyses (t-tests and MTR values distributions) presented in figure 3B, with the frontal regions showing the lowest myelination levels in the first group and a significant increase in myelin content between the first and second group (p < 0.0001). Differently, motor, sensory and occipital cortices show a similar degeneration in cortical myelin content, with significant decreases in MTR values as age increases (all contrasts show p < 0.0001). The premotor cortex confirms its different behavior, with rather stable MTR levels with age and no significant differences in MTR levels between the first and second group.

**Figure 3:**
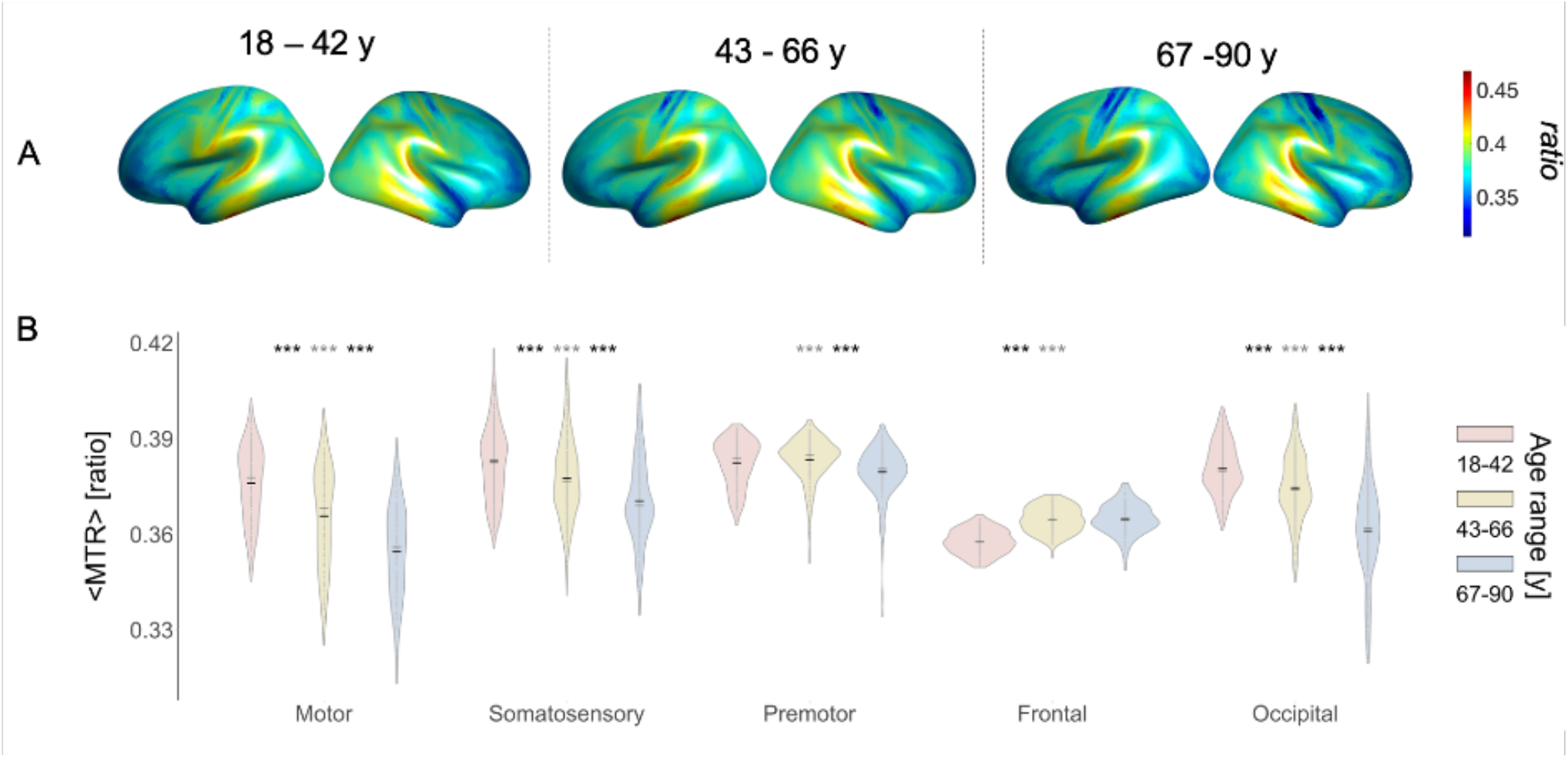
Results for average MTR values in the three age groups; A) MTR average maps for the 7117 isorois for the three age groups separately; B) MTR values distribution for each group in each brain sector of interest (mean in black, median in grey); significance asterisks summarize the results of a t-test between the three age groups for each brain sector separately (*** for p< 0.0001; ** for p < 0.001; * for p< 0.05). Grey asterisks refer to the comparison between the first and the third group.

To better appreciate this age-dependent change in MTR, in Figure 4A.1 the MTR slope is reported as the group-wise result of the linear fit between MTR and age. In the first group there is an increase in cortical myelin content of frontal associative areas, while in the second group frontal areas appear stable (more grey, non-significant regions) and posterior occipital regions show a myelin decrease. In the third group, the anterior areas show a steeper negative slope compared to the second group, while the somatosensory cortex shows some gain in cortical myelin content. The sensory myelin content change with age is different from the rest of the cortex. These observations are confirmed by the quantitative analyses shown in Figure 4B.1: here sensory and occipital regions behave differently from the rest of the cortex with a U-shape trajectory in MTR slope. Especially in the somatosensory regions the myelin content (as assessed with MTR) is positive in the first group (MTR increases), it becomes negative in the second group (MTR decreases) and then it significantly slows down in the third age group (all p-values < 0.0001).

**Figure 4:**
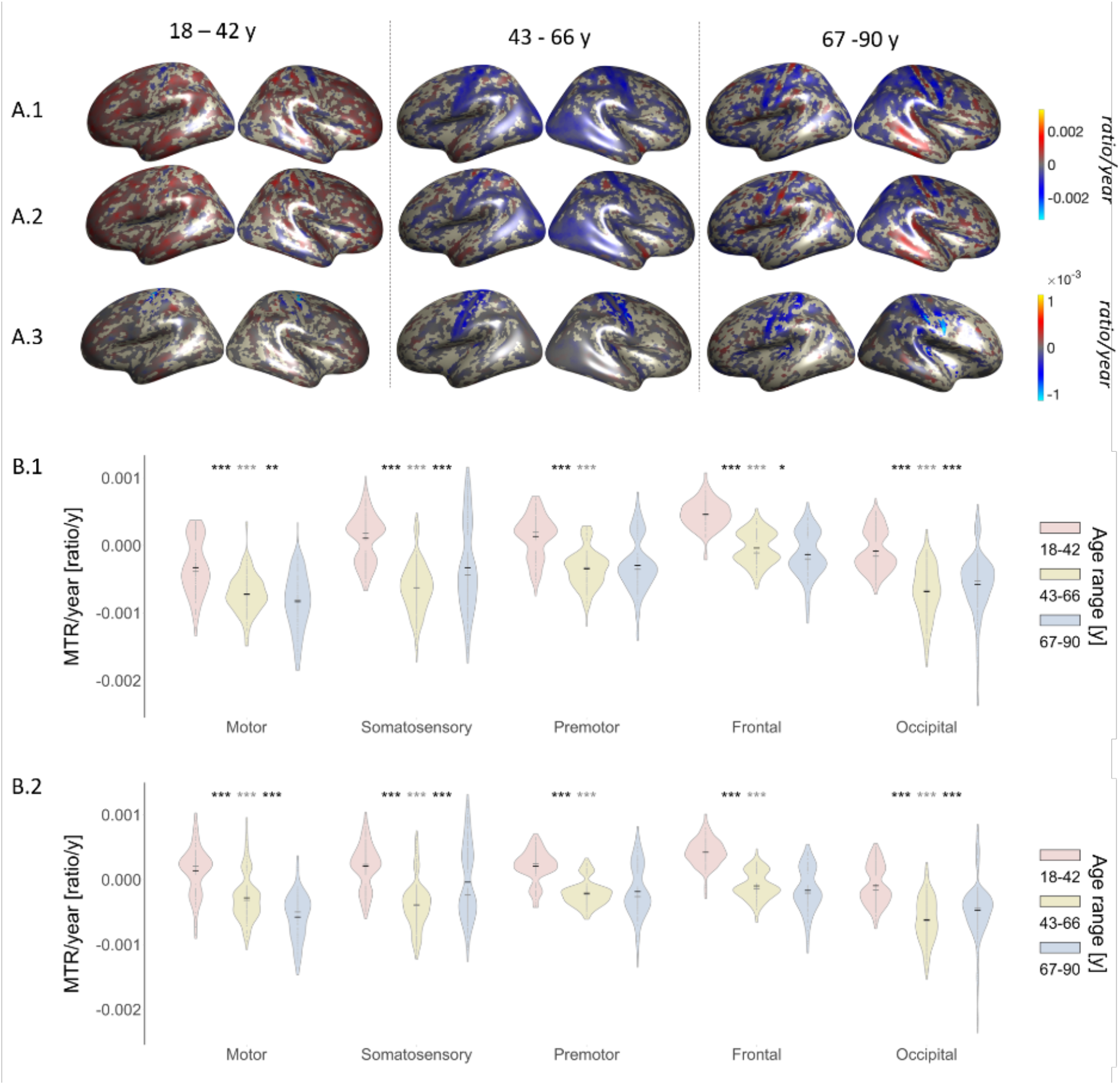
Results for the linear fit between MTR and Age (MTR Slope) in the three age groups (with and without CT ratio regressed out from MTR before computing the linear fit between MTR and Age); A.1) Source map of the slope values from the linear fit between MTR and Age; A.2) Source maps of the slope values from the linear fit between MTR and Age, with CT regressed out as a confound; A.3) Source map of the difference between A. 1 and A.2: the difference is computed only between the values that are significant in both original contrasts; B.1) MTR slope values distribution for each group in each brain sector of interest (mean in black, median in grey); B.2) MTR slope values distribution with CT regressed out as a confound for each group in each brain sector of interest (mean in black, median in grey); significance asterisks summarize the results of a t-test between the three age groups for each brain sector separately (*** for p< 0.0001; ** for p < 0.001; * for p< 0.05). Grey asterisks refer to the comparison between first and third group.

To disentangle the effects of CT on the MTR changes with age, the linear fit in figure 4A.2 was also computed. In this figure, the CT is regressed out from the MTR and then the MTR fit with age (MTR slope) is computed. The MTR slope changes with age shown in figure 4A.1 are not different from the ones shown in Figure 4A.2, suggesting that the MTR slope effects are not due to confounding effects by CT, but rather predominantly determined by actual age-related effects on cortical myelin content. In fact, in figure 4A.3 the differences computed between the significant results in figure 4A.1 and 4A.2 are shown with an extremely small scale compared to MTR changes in both figure 4A.1 and 4A.2. Thus, despite some effects of CT on the MTR visible over parietal areas in the second and third group, the influence of CT over MTR myelin estimate is small compared to the ones really due to age-related effects on the MTR proxy of cortical myelin. These results are confirmed by the quantitative analyses computed over the different sectors of interest, since the patterns observed in figure 4B.1 (t-tests and distributions for MTR slope) and 4B.2 (t-tests and distributions for MTR slope when CT is regressed out) are not different. By comparing figure 4B.1 with figure 4B.2, the only two appreciable differences are: 1) the MTR slope becomes positive in motor cortex in the first group when the CT is regressed out, suggesting that the cortical myelin slope (MTR) is positive in the first group when the effect of CT is not taken into account; 2), a loss of difference between the second and third group in frontal regions, while this difference was significant in the direct linear fit between MTR and age without CT regressions (p <0.05). In all other age groups and in all sectors, the CT regression from the MTR in its fit with age does not change the MTR slope.

As for the results on the age-related changes over myelin content as assessed with the T1w/T2w ratio, in Suppl. Figure 2 and Suppl. Figure 3 the average and slope of T1w/T2w are reported separately for each age group. From a comparison between results in figure 3 and 4 (MTR) and Suppl. Figure 2 and 3 (T1w/T2w ratio), it is clear that myelin estimates computed with MTR and T1w/T2w ratio return different results both in terms of cortical patterns of average myelin content and myelin slope with age. In fact, Suppl. Figure 2A and Suppl. Figure 2B show no remarkable changes between the different age groups in terms of average amount of cortical myelin content when this is assessed with the T1w/T2w ratio proxy. The sensory, premotor and motor regions appear to be the most myelinated in the first age group, which is not far from the results obtained on the MTR average values for the first group (see Figure 3A for a comparison). However, the T1w/T2w ratio does not show the age-related average loss of myelin content which is reported when this is MTR-assessed. In this regard, Suppl. Figure 2B shows numerous non-significant results in the between groups t-test comparisons. Therefore, the T1w/T2w ratio is not particularly sensitive to age-related changes in cortical myelin content. For example, T1w/T2w ratio averages (Suppl. Fig. 2) show no significant changes in myelin content across the three age groups in occipital and premotor areas. Also, the only significant difference in T1w/T2w average values in motor and sensory areas is between the first and third group (p < 0.0001 and p < 0.001, respectively). This latter result suggests that T1w/T2w ratio is more affected by variability compared to the MTR. Finally, a pattern similar to the average MTR-assessed myelin one is visible in frontal regions, since the T1w/T2w ratio average values in frontal regions appear to significantly increase with age (p < 0.0001).

The T1w/T2w ratio slope results reported in Suppl. Figure 3A.1 show very unstable patterns, making it quite difficult to detect clear regional patterns of myelin slope with age. In this figure, the T1w/T2w slope shows a change in myelin content with age. However, apart from a loss of myelin content in the motor areas of the second group, other regions are not well distinguishable in terms of peculiar age-related myelin trajectories. The instability of the T1w/T2w ratio slope is also evident in Suppl. Figure 3A.2, where the CT is regressed out from the T1w/T2w ratio before computing the linear fit with age. These unclear age-related effects on the T1w/T2w ratio are also visible in Suppl. Figure 3A.3, where the only slightly appreciable effect is a contribution of the CT over the T1w/T2w estimate of parietal areas in the second group. In fact, in Suppl. Figure 3B.1 only occipital regions show a T1w/T2w slope which could resemble the pattern observed in the MTR slope (Fig. 4), with a steeper T1w/T2w slope as age increases (p < 0.0001 for all the three contrasts). Differently, the T1w/T2w slope over frontal cortices shows a pattern completely different from the one observed with the MTR: an increase in myelin content with age significantly different between first and second age group (p < 0.0001) is detected. In the premotor, motor and sensory sectors, the T1w/T2w slope shows again less sensitivity compared to MTR at detecting age-related changes in cortical myelin content (less significant differences between age-groups). However, the pattern evidenced in these regions with MTR is reflected in the T1w/T2w ratio slope results. Furthermore, in Suppl. Figure 3B.2 we report results for the between age-groups t-tests and distributions of estimated myelin in the linear fit with age when CT is regressed out from T1w/T2w ratio. From this analysis, it can be seen that T1w/T2w ratio fit with age suffers more from CT regression than from MTR slope. In fact, the t-test between the first and second group on the T1w/T2w slope in the motor cortex is not significant when CT is regressed out from T1w/T2w in its fit with age, while in Suppl. Figure 3B.1 we found a significant result (p < 0.05). Also, in the premotor cortex the first and third group show a significant difference in Suppl. Figure 3B.2 (T1w/T2w slope when CT is regressed out from T1w/T2w ratio), while this was not the case in Suppl. Figure 3B1 (T1w/T2w slope when CT is not regressed out from T1w/T2w ratio). Finally, no significant difference is also detected in the occipital cortex in T1w/T2w slope between the second and third group when the CT is regressed out from T1w/T2w ratio, while this contrast was significant (p<0.0001) in Suppl. Figure 3B.1. Importantly, the peculiar tendency toward a U-shaped trajectory of sensory and occipital regions is detected also with the T1w/T2w ratio.

Due to different results obtained using the MTR and T1w/T2w proxy, we also computed the linear fits shown in figure 5, in Suppl. Figure 4 and Suppl. Figure 5. In Figure 5A, the linear fit between CT and MTR is reported, which shows a positive linear relationship between the MTR and CT over parietal areas but a negative linear relationship between the MTR and CT in the rest of the brain regions, especially in the first age group. These results are confirmed by the between age-groups t-tests in Figure 5B, where the parietal district shows a positive linear relationship between CT and MTR which tends to increase with age. The occipital and frontal regions show a negative relationship between CT and MTR in the first age group. However, the relationship between CT and MTR in these areas becomes significantly positive with age, with frontal areas showing a stronger positive relationship between the two metrics across all age groups, while the occipital regions show this difference only between the first and the second group (p < 0.0001).

**Figure 5:**
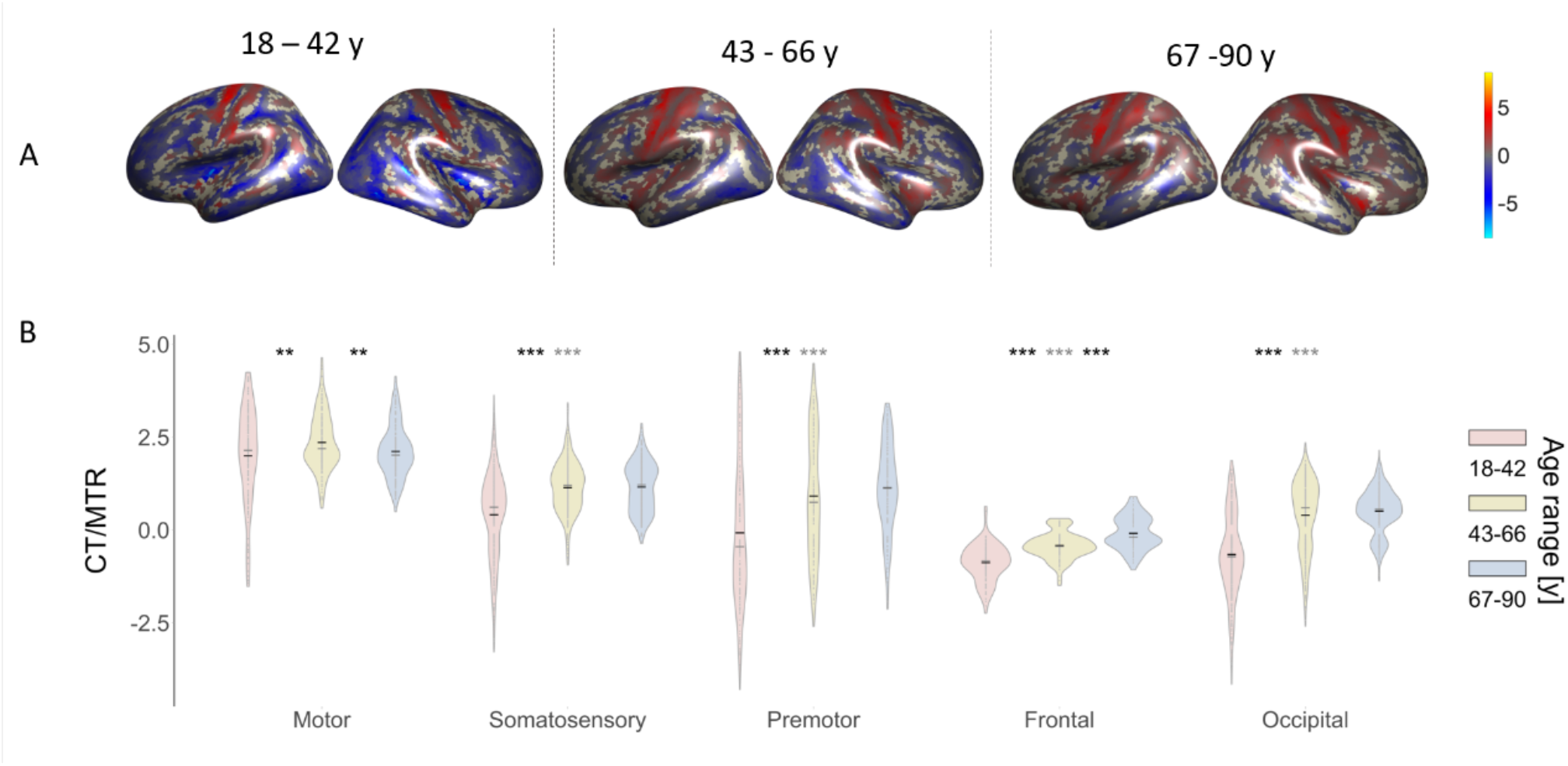
Results for the linear fit between CT and MTR in the three age groups; A) Source map of slope values from the linear fit between CT and MTR; B) Slope distribution from the linear fit between CT and MTR for each group in each brain sector of interest (mean in black, median in grey); significance asterisks summarize the results of a t-test between the three age groups for each brain sector separately (*** for p< 0.0001; ** for p < 0.001; * for p< 0.05). Grey asterisks refer to the comparison between the first and the third group.

In Suppl. Figure 4A, the same linear fit is computed between the CT and T1w/T2w proxy of cortical myelin. This fit shows similar results as in Figure 5A, especially in first and second age groups, but with inconsistent patterns in sensorimotor areas. By comparing the results in the third group in Figure 5A and Suppl. Figure 4A, it can be also noticed that while a decrease in CT in every brain region (with the exception of the parietal ones) determines an increase in the T1w/T2w estimate (negative relationship), this is not true for the CT vs MTR fit. These results are confirmed by the results reported in Suppl. Figure 4B where the between-groups t-tests and distributions are shown. In Suppl. Figure 4B, in fact, the relationship between CT and T1w/T2w is negative especially in frontal, premotor and occipital regions until the last age group but this is not true for the fit between CT and MTR which shows a progressive tendency with age towards a positive relationship between CT and myelin.

Given the differences evidenced so far between MTR and T1w/T2w, the last fit was computed as shown in Suppl. Figure 5A, representing the direct linear fit between the MTR and T1w/T2w ratio. Here the two cortical myelin proxies do not show a uniform relationship with each other as expected. In fact, their relationship changes depending on the brain region taken into account. Suppl. Figure 5B, shows the results for the between-groups t-tests and distributions of the fit between T1w/T2w and MTR. The two proxies show a positive relationship that tends to increase with age ubiquitously on cortex with the exception of the sensory area.

## Discussion

### Age-related changes in Cortical Thickness

The goal of this work is to provide the reader with an investigation of age-related brain structural changes on a large, single-site and homogeneously distributed cohort of participants aged between 18 and 89 years. We focussed on optimally exploiting age related degeneration patterns by reducing sample variability. In particular we addressed: 1 regional specific effects of aging on cortical thickness; 2. Regional specific effects of aging on cortical myelin concentration; 3. differences between the T1w/T2w ratio and MTR proxies in estimating cortical myelination.

It has been shown here that aging processes not only affect the grey matter thickness, but also myelin concentration. Importantly, this happens differently depending on the region of the cortex. Moreover, a general loss of cortical thickness can be appreciated with age in all brain regions, with frontal and premotor cortex showing the highest thickness values (Figure 1A and1B). The distinct behaviours of different cortical districts appear evident in the slope of CT with age, which describes the rate at which thinning occurs. In fact, CT slope shows premotor and motor cortex to be very different compared to the rest of the brain in terms of thinning rate across the different age-groups. In Figures 2A.1 and 2B.1, the premotor and motor cortices show in fact a faster thinning in the first group and then a slow-down in thinning rate with age. Differently, the sensory cortex shows a rather constant thinning over the life span, with a stabilization in the second group which does not differ from thinning rate in the third age group. The frontal and occipital cortices also show a peculiar pattern, with thinning rate getting faster with age. Our results on the CT slope are partially in line with previous results (e.g. Salat et al., 2004; Thambisetty et al., 2010; Fjell et al., 2009). For instance, also Salat and colleagues (2004) found the fastest thinning rate by middle-age to occur in the motor cortex, while Thambisetty and colleagues found an anterior-posterior gradient in the decline rates, with both frontal and parietal areas showing the largest thinning. However, it must be noticed that our age sample ranges from 18 to almost 90 years, with a large cohort of participants (N = 610) included in our sample. Instead, Thambisetty et al.’ work (2010) only included participants aged from 60 years old and Salat et al. (2004) included only 106 participants for the same life span investigated in our work. Therefore, our finding of a fast thinning in motor areas already in the first group of age depends on the fact that, on one hand Thambisetty and colleagues did not investigate thinning before 60 years of age, while, on the other hand, Salat and colleagues included a sample size that might have been too small to appreciate small gradual changes. Differently from our results and aforementioned previous works, McGinnins and colleagues (2011) found that sensory and motor cortices show highest thinning predominantly in the late part of life, partially overlapping with our third age group. However, the number of participants included in McGinnis et al. (2011)’ age groups was smaller than ours (mean N per age-group = 79; N in older age group = 38). Therefore, while their analysis was sensitive to the general larger thinning in the motor cortex compared to the rest of the brain (effect that we also found), their ANOVA tests in the between groups analyses might have suffered from the different sample sizes in the compared groups, leading to non-significant differences in the comparisons of CT slope in the motor cortex until their last group of age. Hence, while our evidence converges with some of the previous works, we were additionally able to extract finer differences between age groups. Most importantly, none of the previous works highlighted the difference in aging patterns of motor vs. sensory areas. In fact, sensory regions show a slower thinning compared to premotor and motor cortices in the first age group, with a significant increase in the thinning rate between the first and second age groups and a final thinning stabilization between the second and third age group, therefore from middle-age on. On the contrary, the premotor and motor regions show a fast thinning rate at first which gradually slows down with age. It is worth to notice that these differences cannot be ascribed to field inhomogeneities in the MR head coil, since both motor and sensory areas lie almost in the same coil area during the MRI scan. The similarity between premotor and motor cortices and their dissimilarity with the sensory cortex are also evident in the CT slope when the MTR is regressed out in figure 2A.2 and 2B.2. In fact, when CT (with MTR regressed out) is fitted against age, sensory cortices show a stable thinning over life (no differences between age groups), while premotor and motor regions still show a larger thinning compared to sensory cortices and a slowdown in thinning with age. This dissociation between sensory and premotor/motor cortices reminds the results of a recent work by Wagstyl and colleagues (2020) which shows by means of a quantitative 3D laminar atlas of the cerebral cortex, layers III, V and VI to exhibit opposite behaviours in the premotor and motor cortex compared to the sensory cortex. Although Wagstyl et al. (2020) did not investigate these different regional-dependent behaviours in individuals with different ages, their results support the dissociation we found in the aging patterns, in terms of cortical thickness, of these two sectors of the human brain.

### Age-related changes in Cortical Myelin

Age-related cortical myelin changes were assessed using two different proxies of myelin content, namely the MTR and T1w/T2w ratio. Both MTR and T1w/T2w results show premotor, motor and sensory regions as the most myelinated districts of the cortex, in accordance with previous studies on cortical myelin content in humans (e.g., Patel et al., 2020; Shafee et al., 2015; Glasser and Van Essen., 2011). These results were consistent in both MTR and T1w/T2w cortical myelin averages estimates (Figure 3 and Suppl. Figure 2).

### Age-related changes in Cortical Myelin as assessed with MTR

MTR slope shows a loss of cortical myelin content with age in all brain regions (Figure 4). Also in these myelin estimates, different patterns were found in the myelin thinning rate depending on the examined areas. In fact, we found a progressive increase in the rate of myelin loss in anterior regions (frontal, motor and premotor), while sensory and occipital regions showed a U-shaped trajectory (Figure 4B.1). It is worth noticing that these patterns showing a progressive age-related increase in myelin loss in anterior regions and U-shaped trajectory in posterior ones are even more pronounced when the effect of CT is regressed out from the linear fit between MTR and age (Figure 4B.2). This suggests that such patterns are mainly due to myelin and not to the confounding effects of CT on cortical myelin content as assessed with MTR. Our results on the average amount of cortical myelin (as assessed with the MTR proxy: Figures 3A and 3B) are in line with Patel et al. (2020) and Wu et al. (2016), who used the MTR to assess cortical myelin content. However, our results on the myelin thinning rate do not confirm Wu and colleagues’ findings as they could not detect any significant age-related change in cortical myelin content in sensory and motor areas. Their nil finding is probably due to the small sample size employed, as they included 66 participants to investigate MTR changes over a huge life span (30 to 85 years). Differently, Karolis et al. (2019) also employed the MTR to assess cortical myelin loss with age in a cohort of 97 participants aged between 20 and 74 years. They found age-related changes in myelin content in parietal regions, as we report in our results. However, none of the previous works reported the dissociation between sensory and motor regions, a dissociation that we also found in the age-related changes of the MTR estimates (with and without CT regression). In fact, sensory and motor regions show different age-related trajectories also in myelin concentration, and not only in cortical thinning. Moreover, common age-related changes in sensory and occipital cortices can be noticed in Figure 4B.1 and 4B.2 as opposed to the different age-related changes in more anterior regions (frontal, premotor and motor). These commonalities between somatosensory and occipital regions were also found in the investigation of the laminar features of the different brain regions by Wagstyl and colleagues (2020). In fact, they found common structural properties between sensory and occipital regions as opposed to the ones typical of frontal, premotor and motor districts.

### Age-related changes in Cortical Myelin as assessed with T1w/T2w ratio

Furthermore, we found that the age-related changes in myelin content assessed with the T1w/T2w proxy partly differ from the ones obtained with MTR. Frontal, premotor and motor myelin thinning with age assessed with T1w/T2w ratio resembles an inverted U-shaped trajectory, as (Suppl. Figure 3B1). In fact, frontal and premotor cortex show a faster thinning in the first group which slows down and stabilizes in the second and third age groups. The motor cortex shows a clear inverted U-shaped trajectory, with significant decrease in myelin thinning in the second compared to the first age group, and a late acceleration in the third age group. These results are in accordance with the work by Grydeland and colleagues (2013) who also employed T1w/T2w ratio as a myelin proxy to investigate age-related myelin changes over the life span and found an inverted U-shaped trajectory in anterior regions. However, our results differ from Grydeland and colleagues’ as we find a dissociation between age-related changes in sensory regions vs anterior regions also when myelin was assessed with the T1w/T2w ratio. In fact, we found occipital and somatosensory cortices to show a progressive increase in myelin thinning rate with age, with a tendency towards a stabilization in myelin thinning in the second age group.

### T1w/T2w ratio is more affected by Cortical Thickness compared to MTR

It should be noticed that the CT regression (from the fit between the T1w/T2w and age) affects more the results in T1w/T2w ratio estimates compared to the regression of CT from the fit between the MTR and age (which only led to an enhancement of the effects seen in the relationship between MTR and age). In fact, comparing Suppl. Figure 3B.1 and Suppl. Figure 3B.2, in premotor, motor and occipital results change in the relationship between T1w/T2w and age when CT is regressed out from the T1w/T2w before computing the linear fit with age. In particular, CT regression from the T1w/T2w determines a more pronounced difference in premotor regions between the first and second group, therefore leading to a more significant deceleration in myelin thinning with age in this region. In the motor cortex, changes in T1w/T2w slope due to the CT regression are detected in the first and second group, which do not differ in thinning rates like they did when CT regression was not implemented. Also, the significant thinning increase within the occipital cortex in the third vs. second group disappears, suggesting that the CT regression from T1w/T2w leads to estimates similar to the ones obtained with the MTR proxy. Therefore, we show that the T1w/T2w ratio is relevantly more susceptible to the CT regression than the MTR estimate. In other words, T1w/T2w myelin estimates are more affected by values of cortical thickness at each surface location than MTR estimates. In this regard, it has to be considered that the T1w/T2w ratio depends on the intensity of the T1-weighted and T2 weighted images. At the same time, the CT extraction depends on the intensity of the T1-weighted image as CT is ultimately obtained from the contrast differences between grey matter and white matter, as evaluated during the segmentation process. Conversely, the MTR is obtained from the ratio between two images acquired with a specifically designed sequence which is different from the T1-weighted image used for the T1w/T2w computation and CT extraction. Actually, the MTR pipeline only involves the original T1-weighted image for rigid-body co-registration. For these reasons, the greater co-dependency between the T1w/T2w ratio and CT can be explained with a shared dependency of the two metrics on the intensity of the common T1w image. Therefore, we consider the results obtained with the MTR (which is also less affected than the T1w/T2w contrast by iron molecules accumulation (Pareto et al., 2020)) as more stable and closely reflecting changes in cortical myelin changes. In this light, in Figure 5 and Suppl. Figure 4, the relationship between CT and MTR and CT and T1w/T2w were respectively computed. Due to the strong impact of CT removal on T1w/T2w fit with age previously discussed, the results in Figure 5A and 5B should be considered a more reliable description of the relationship between myelin concentration and CT compared to Suppl. Figure 4A and 4B. The direct fit between CT and MTR suggests that in premotor, motor and sensory areas, the greater the thickness, the greater the myelin. These results are also confirmed by the linear fit between CT and myelin content as assessed with the T1w/T2w ratio in Suppl. Figure 4A and 4B, with the exception of sensory cortex where the relationship between CT and T1w/T2w ratio is non-existent in all three age groups. These results are in line with previous works correlating T1w/T2w ratio maps with cortical thickness maps (Shafee et al., 2015; Glasser et al., 2011). However, none of the previous works explored the changes in the relationship between CT and cortical myelin content with age, as we did in this work. Therefore, for the first time our results provide information on the relationship between CT and cortical myelin content with age. We show that during young adulthood, thickness is positively correlated with myelin only in sensory and motor areas, while this relationship becomes a feature of the whole cortex with further aging.

### Relationship between T1w/T2w ratio and MTR

Finally, we were interested in the relationship between the T1w/T2w and MTR proxy of myelin from a methodological point of view. Building on previous results reporting differences between the two proxies of cortical myelin content, we further assessed whether between the two metrics there was a uniform relationship across the brain, as it would be theoretically expected. We found that the two metrics show an unstable and non-uniform relationship across brain regions, as it can be appreciated in Suppl. Figure 5. This result is consistent with previous works showing that the MTR and T1w/T2w ratio show only moderate correlations, with the T1w/T2w ratio being more affected by iron molecules than MTR (e.g. Nakamura et al., 2017; Pareto et al., 2020). To our knowledge, this is the first work to provide a direct linear comparison across the whole cortex between the two proxies’ over the life span. Thanks to this analysis, we show that the T1w/T2w and MTR share a positive relationship all over the brain with the exception of temporal cortices. This relationship between the two proxies tends to increase with age especially in frontal, motor and occipital regions, while in premotor and sensory cortices it does not appear affected by age.

To summarize our results, we report that the largest cortical thickness values are found in more anterior regions (frontal and premotor), followed by motor, sensory and occipital regions. Therefore, we found an anterior-posterior gradient of average cortical thickness values. We did not find the same gradient when cortical thinning was examined, despite previous works supporting this framework (Thambisetty et al., 2010). We show here that cortical thinning regional patterns are more complex than merely bound to a law of spatial gradient degeneration. Specifically, we found frontal and occipital areas to increase in their thinning rate with age, while motor and premotor areas show a greater thinning rate in the first age group which slows down later with age. Sensory areas behave differently, as their thinning rate is more stable over life-span (see Figure 2). In this regard, the work by Thambisetty and colleagues (2010) investigated thinning in a group of 66 adults longitudinally followed up from their 60s up to their 80s. Despite the great reduction of sample variability due to the longitudinal study, the age-related thinning did not cover extensively the entire life-span. Only the increased thinning in frontal areas with age and the slowdown in thinning of more posterior areas were reported. For this reason, parietal suffering from a faster thinning rate compared to frontal regions in early adulthood could not be appreciated as we did here.

The results on the cortical myelin concentration show that MTR is a more reliable proxy for cortical myelin content as its results are less affected by CT regression than the T1w/T2w ratio’s. In general, we find the most myelinated are in the parietal lobe, with a myelin concentration peak in the somatosensory cortices. Differently, frontal regions show the least average amount in myelin content (Figure 3). These results are in line with previous findings (e.g. Shafee et al., 2015). However, we are to our knowledge the first to report a dissociation in the rate of age-related cortical myelin changes between anterior and posterior areas and especially between sensory and motor areas. In fact, we report that motor regions increase their myelin loss rate with age, while somatosensory areas show a U-shaped trajectory, similar to the one observed in occipital areas. This structural dissociation between somatosensory and motor areas, found both in cortical thinning and cortical myelin age-related changes, was until present only reported in works investigating the structure of the brain at the laminar level (Wugstyl et al., 2020). Our capability to detect these differences between the two sectors in vivo is to be ascribed to the large sample employed in our study (N = 610) and the comparable sample sizes included in the three investigated age groups (N ~ 200 in each age group).

It is here then proposed that the anterior-posterior gradient hypothesis might be too simplistic to account for the regional differences in brain degeneration. First, the anterior-posterior gradient hypothesis was formulated on the basis of results obtained on samples representative of short age span. Secondly, this account was formulated on the basis of results only obtained on grey matter degeneration, without taking into account cortical myelin concentration (Thambisetty et al., 2010; Resnick et al., 2003). The same issue can affect the last in first out account (Raz, 2000). In fact, a complete picture describing age-related changes in the brain should take into account both cortical thickness and cortical myelin as well as rely on results from studies including large samples representative of sufficiently long age-spans.

## Conclusions

In the present work we used structural MRI recordings from a big cohort of participants (N = 610) from the CAM-Can Project to address region specific aging in cortical thickness and myelination. To our knowledge, this is the first work where structural age-related brain changes are investigated employing 1) a large and uniformly age distributed data sample, 2) recordings from a single site, 3) comparable sample sizes in the three age groups and 4) two different proxies (T1w/T2w ratio and MTR) to assess cortical myelin concentration. It has been shown here that cortical thickness and myelin content both degenerate with aging, showing different regional aging patterns. In particular, we provide evidence for a dissociation between sensory and motor cortices, both in terms of cortical and myelin thinning. This has been highlighted only in works investigating the brain structural properties at the laminar level. Interestingly, myelin estimates further show commonalities between somatosensory and occipital cortices as opposed to the frontal, premotor and motor age-related degeneration trajectories. This dissociation is particularly evident when age-related brain changes are investigated using the MTR, while the T1w/T2w ratio is here shown to be more affected by cortical thickness and therefore to be a less reliable proxy of cortical myelin content.

## Authors’ contributions

Conceptualization: AB., DT., PB.; Methodology: AB, DT; Software: AB, DT; Formal analysis: AB, DT; Resources: AB, DT; Data Curation: AB, DT; Writing - Original Draft: AB; Writing - Review & Editing: AB, DT, PB; Visualization: DT; Supervision: DT, PB;

## Declaration of interest

The authors declare no conflict of interests.

## Data Availability Statement

Data analyzed in this study are part of the Cambridge Centre for Aging and Neuroscience public dataset (Cam-CAN; Shafto et al., 2014; Taylor et al., 2017) and are publicly available.

## Aknowledgments

Data collection and sharing for this project was provided by the Cambridge Centre for Ageing and Neuroscience (CamCAN). CamCAN funding was provided by the UK Biotechnology and Biological Sciences Research Council (grant number BB/H008217/1), together with support from the UK Medical Research Council and University of Cambridge, UK.

## Notes

https://www.cam-can.org/index.php?content=dataset

### Competing Interest Statement

The authors have declared no competing interest.

