## Supplementary material for "Region-specific impact of aging on cortical myelination and thickness"

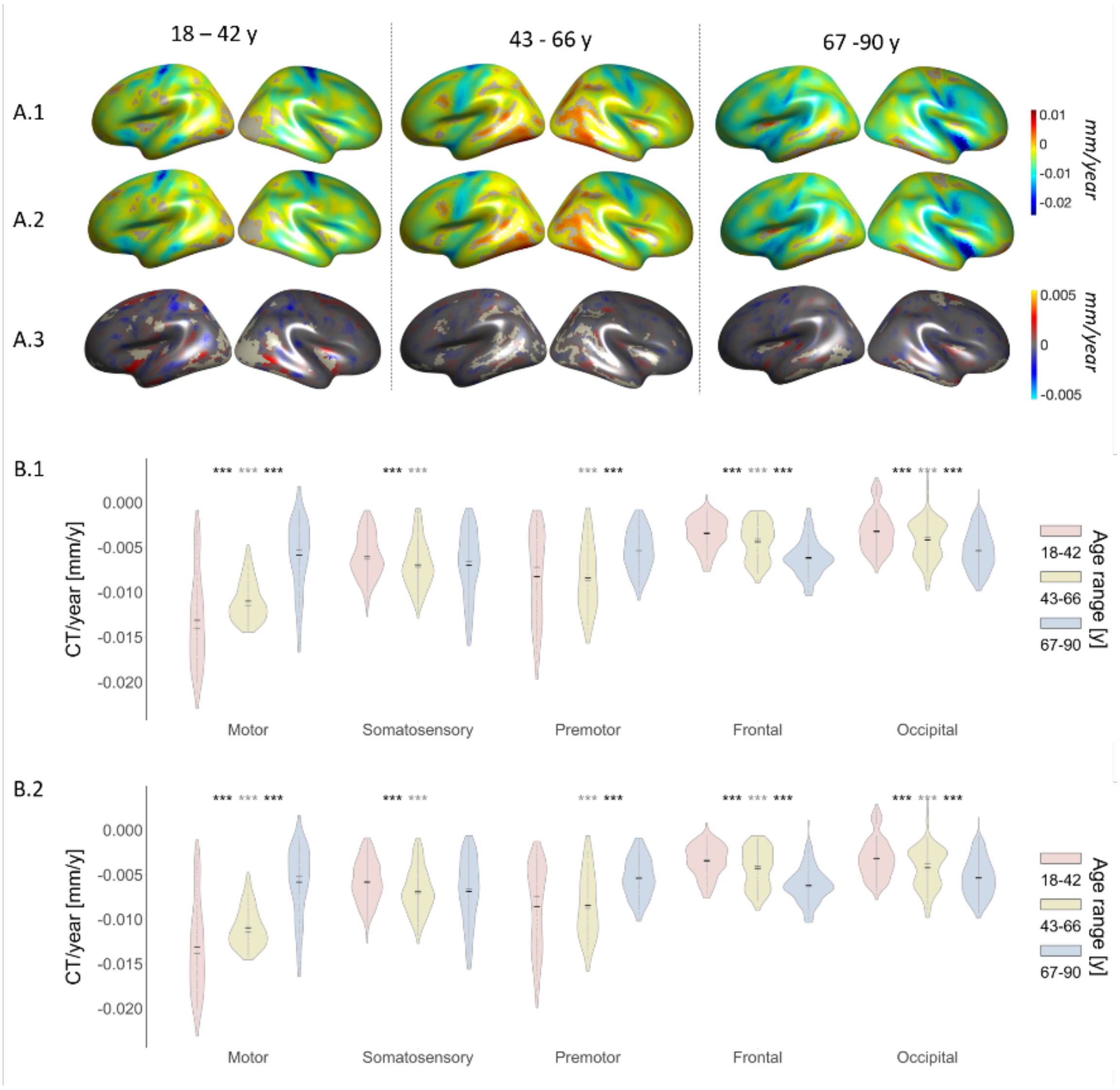


**Suppl. Figure 1**: Results for the linear fit between Cortical Thickness and Age (Cortical thinning/CT Slope) in the three age groups (with and without T1w/T2w ratio regressed out from CT before computing the linear fit between CT and Age); A.1) Source map of the slope values from the linear fit between CT and Age; A.2) Source maps of the slope values from the linear fit between CT and Age, with T1w/T2w ratio regressed out as a confound.; A.3) Source map of the difference between A.1 and A.2: the difference is computed only between the values that are significant in both original contrasts; B.1) CT slope values distribution for each group in each brain sector of interest (mean in black, median in grey); B.2) CT slope values distribution with T1w/T2w ratio regressed out as a confound for each group in each brain sector of interest (mean in black, median in grey); significance asterisks summarize the results of a t-test between the three age groups for each brain sector separately (*** for p< 0.0001; ** for p < 0.001; * for p< 0.05). Grey asterisks refer to the comparison between the first and the third group.

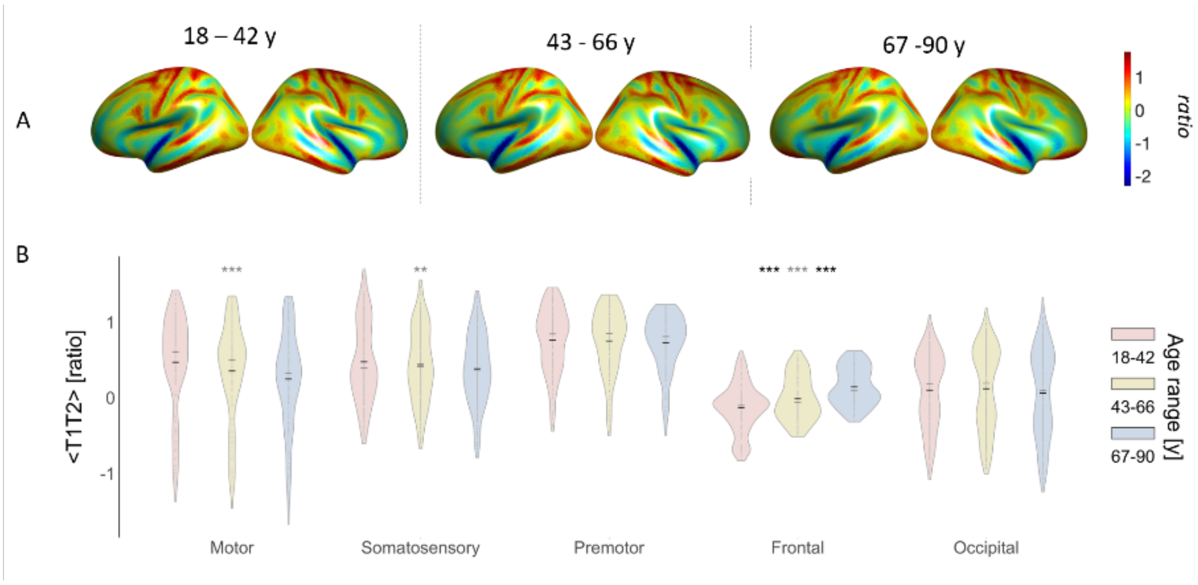


**Suppl. Figure 2**: Results for average T1w/T2w ratio values in the three age groups; A) T1w/T2w ratio average maps for the 7117 isorois for the three age groups separately; B) T1w/T2w ratio values distribution for each group in each brain sector of interest (mean in black, median in grey); significance asterisks summarize the results of a t-test between the three age groups for each brain sector separately (*** for $p< 0.0001$; ** for $p < 0.001$; * for $p< 0.05$). Grey asterisks refer to the comparison between the first and the third group.

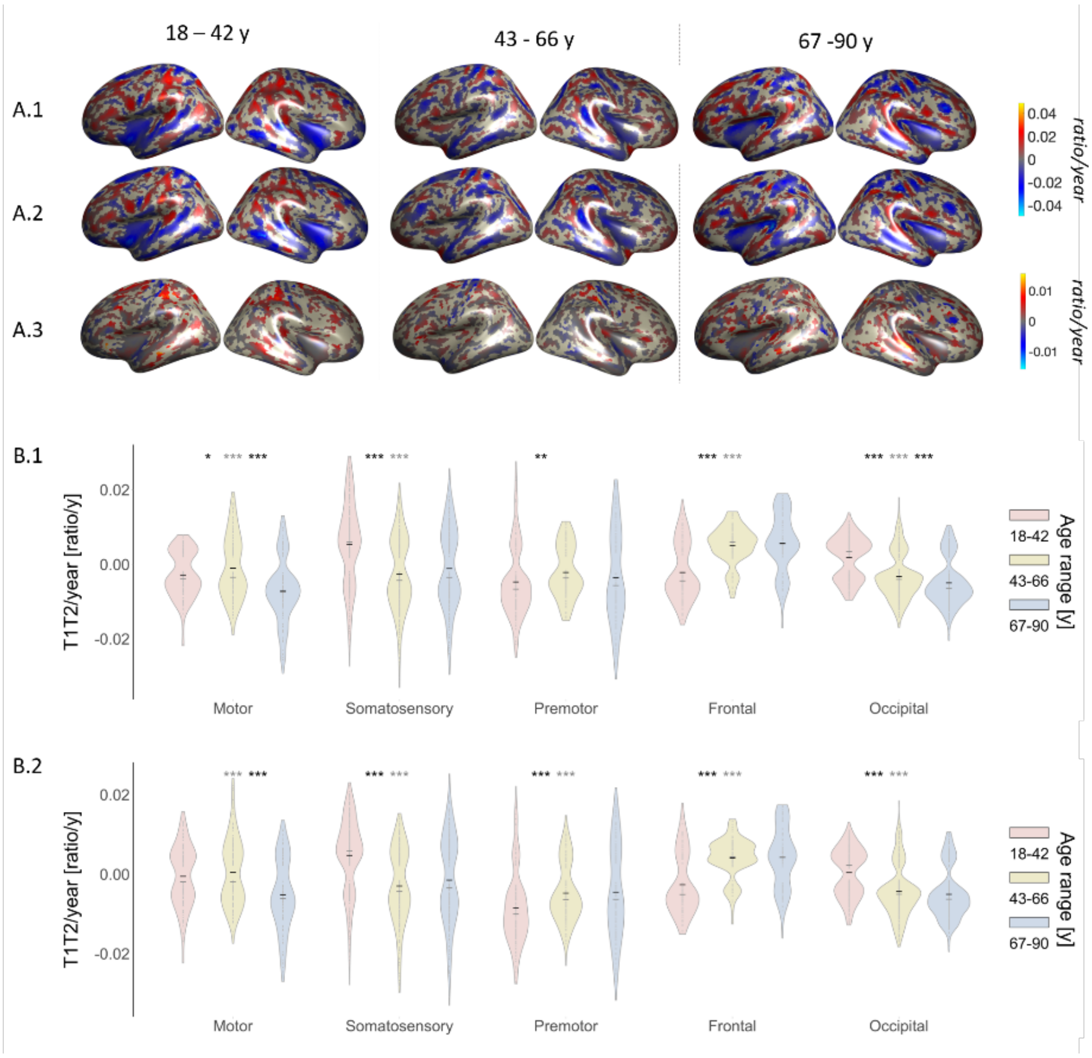


**Suppl. Figure 3**: Results for the linear fit between T1w/T2w ratio and Age (T1w/T2w ratio Slope) in the three age groups (with and without CT ratio regressed out from T1w/T2w ratio before computing the linear fit between T1w/T2w ratio and Age); A.1) Source map of the slope values from the linear fit between T1w/T2w ratio and Age; A.2) Source maps of the slope values from the linear fit between T1w/T2w ratio and Age, with CT regressed out as a confound.; A.3) Source map of the difference between A.1 and A.2: the difference is computed only between the values that are significant in both original contrasts; B.1) T1w/T2w ratio slope values distribution for each group in each brain sector of interest (mean in black, median in grey); B.2) T1w/T2w ratio slope values distribution with CT regressed out as a confound for each group in each brain sector of interest (mean in black, median in grey); significance asterisks summarize the results of a t-test between the three age groups for each brain sector separately (*** for $p < 0.0001$; ** for $p < 0.001$; * for $p < 0.05$). Grey asterisks refer to the comparison between the first and the third group.

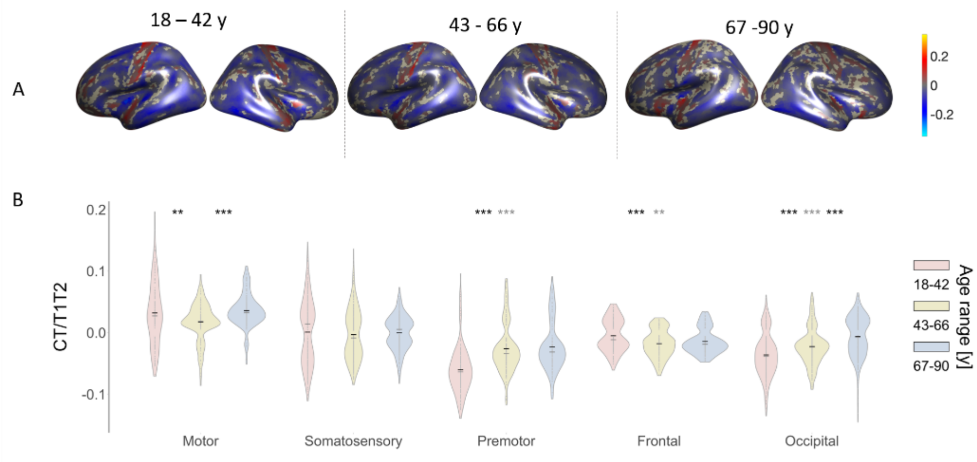


**Suppl. Figure 4**: Results for the linear fit between CT and T1w/T2w ratio in the three age groups; A) Source map of slope values from the linear fit between CT and T1w/T2w ratio; B) Slope distribution from the linear fit between CT and T1w/T2w ratio for each group in each brain sector of interest (mean in black, median in grey); significance asterisks summarize the results of a t-test between the three age groups for each brain sector separately (*** for $p< 0.0001$; ** for $p < 0.001$; * for $p< 0.05$). Gray asterisks refer to the comparison between the first and the third group.

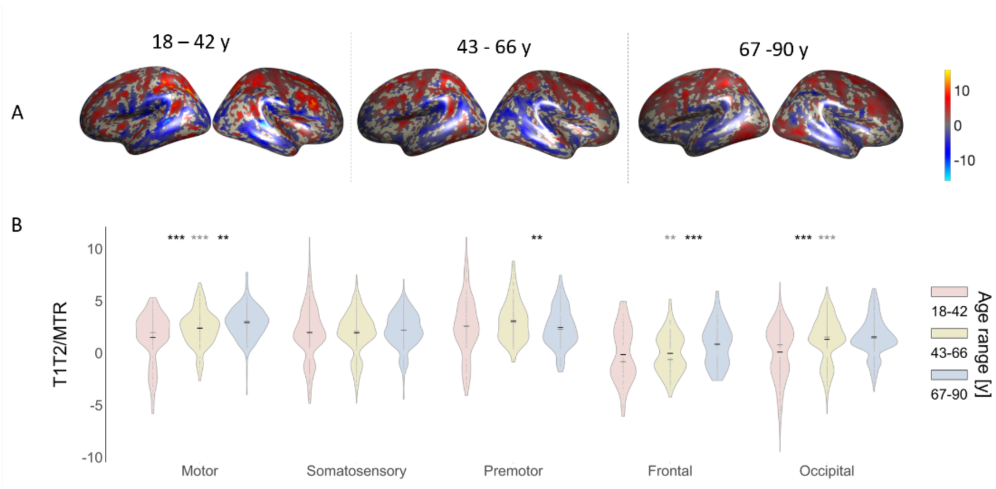


**Suppl. Figure 5**: Results for the linear fit between T1w/T2w ratio and MTR in the three age groups; A) Source map of slope values from the linear fit between T1w/T2w ratio and MTR; B) Slope distribution from the linear fit between T1w/T2w ratio and MTR for each group in each brain sector of interest (mean in black, median in grey); significance asterisks summarize the results of a t-test between the three age groups for each brain sector separately (*** for $p< 0.0001$; ** for $p < 0.001$; * for $p< 0.05$). Gray asterisks refer to the comparison between the first and the third group.

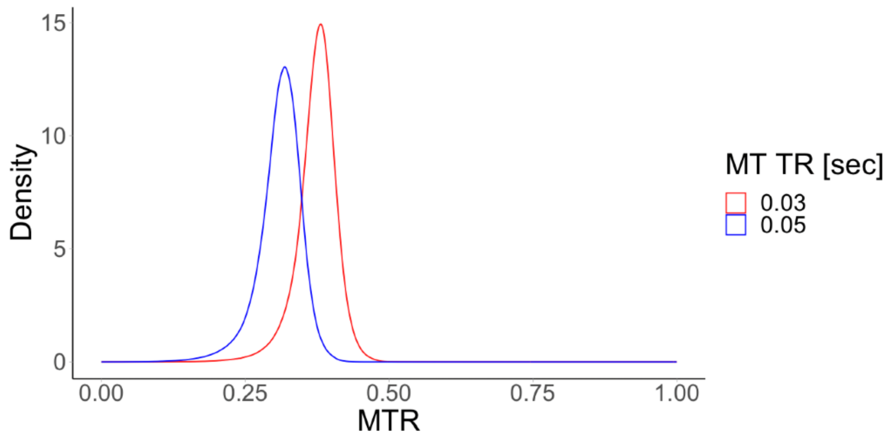


**Suppl. Figure 6**: Density plot for the distribution of MTR values with respect to the MT sequence repetition times for all subjects and all vertices in the selected sectors.

| Study | Sample size | Multisite | Longitudinal | Age-range |
|---|---|---|---|---|
| Fjell et al., 2009 | 883 (avg N = 147) | yes (6 indep. samples) | no | 19 – 81 (avg across samples) |
| Frangou et al., 2020 | 17075 (avg N = 205) | yes (83 indep. samples) | no | 21 – 55 (avg across samples) |
| Hutton et al., 2009 | 48 | no | no | 20 – 60 |
| Lemaitre et al., 2012 | 216 | no | no | 18-87 |
| McGinnis et al., 2011 | 316 | no | no | Young = 18 - 29<br>Middle-age = 30 - 59<br>Old = 60 - 79<br>Old-old = >80 |
| Resnick et al., 2003 | 24 | no | yes (5 years) | 69.5 ± 5.6 at baseline |
| Salat et al., 2004 | 106 | no | no | Young = 18 - 31<br>Middle-age = 41 - 57<br>Old = 60 – 93 |
| Thambisetty et al., 2010 | 66 | no | Yes (8 years) | 60 - 84 at baseline |

**Suppl. Table 1:** List of studies investigating cortical thinning and grey matter changes with age. Sample size is reported for each study. When the study included more than one sample, "avg N" in brackets refer to the average number of participants across samples. Multisite specifies whether a study included MRI images acquired with different scanners across participants. Longitudinal refers to whether a study included at least two time points at which participants where scanned (in this case, the years range is reported). Age-range refers to the age-range investigated in the study, with average age-ranges reported for studies including more than one sample.

| Study | N Total | Multisite | Longitudinal | Age-range | Proxy |
|---|---|---|---|---|---|
| Deoni et al., 2015 | 215 | no | no | 1 – 6 | T1w/T2w ratio |
| Grydeland et al., 2013 | 339 | no | no | Young = 8.4 - 19.7<br>Old = 19.7 - 83.1 | T1w/T2w ratio |
| Karolis et al., 2019 | 97 | no | no | Young = 26.4 ± 4.7<br>Middle= 46.6 ± 6.7<br>Old = 65.9 ± 5.2 | MTR |
| Kwon et al., 2020 | 225 | yes | yes | 12 - 21 | T1w/T2w ratio |
| | 686 | no | no | 22 - 35 | |
| Shafee et al., 2015 | 1555 | no | no | 18 - 35 | T1w/T2w ratio |
| Wu et al., 2016 | 66 | no | no | 30 - 85 | MTR |

**Suppl. Table 2:** List of studies investigating cortical myelin changes with age. Sample size is reported for each study. Note that in Kwon et al. (2020) two different samples were studied separately to assess cortical myelin changes in samples representative of different age ranges. Multisite specifies whether a study included MRI images acquired with different scanners across participants). Age-range refers to the age-range investigated in the study. Proxy refers to the metric employed in the study in order to assess cortical myelin concentration.
